# The Role of Terminal Uridyl Transferases in the Circadian Rhythm

**DOI:** 10.1101/2023.03.13.516290

**Authors:** Wei Song, Juan Wang, Weilan Piao, Yuening Yang, Ahyeon Son, Hyeshik Chang, Xianghui Wen, Honglei Zhang, Chong Li, Daxiang Na, Yanming Lu, Jerome Menet, V. Narry Kim, Hua Jin

## Abstract

The 3’ terminal oligo-uridylation, a post-transcriptional mRNA modification, is conserved among eukaryotes and drives mRNA degradation, thereby affecting several key biological processes such as animal development and viral infection. Our TAIL-seq experiment of mouse liver mRNA collected from six zeitgeber times reveals transcripts with rhythmic poly(A) tail lengths and demonstrates that overall 3’ terminal uridylation frequencies at mRNA poly(A) tail very-ends undergo rhythmic change. Consistently, major terminal uridylyl transferases, TUT4 and TUT7, have cycling protein expression in mouse liver corresponding to 3’ terminal uridylation rhythms, indicating that the cycling expression of TUTases correlates with the rhythmic pattern of uridylation. Furthermore, the double knockdown of TUT4 and TUT7 in U2OS cells lengthens the circadian period and decreases the rhythmic amplitude of clock gene expression. Our work thoroughly profiles the dynamic changes in poly(A) tail lengths and terminal modifications and uncovers uridylation as a post-transcriptional modulator in the mammalian circadian clock.

## Introduction

Poly(A) tails universally exist at the ends of most eukaryotic mRNAs and long non-coding RNAs ^1^ and the lengths of poly(A) tails are tightly regulated in both the nucleus and the cytoplasm through various mechanisms ^2^. Some maternal mRNAs with short poly(A) tails are stored, and later activated through elongation of their poly(A) tails in the cytoplasm during oocyte-to-embryo switch ^3-5^. In depth studies of poly(A) tails have elucidated their roles in mRNA stability, transport and translation ability ^6^.

Previous research has found that poly(A) tails possess other ribonucleotides such as uridine or guanosine, and the dynamic processing and modification of poly(A) tails occur due to and mostly depending on the Terminal Nucleotidyl Transferase (TENT) family ^7-12^. These TENTs modify the 3’ ends of mRNAs and regulate their abundance as well as their translation. At present, eleven TENTs have been identified in human. Based on substrate preference for adenosine triphosphate (ATP) or uridine triphosphate (UTP), TENTs can be divided into the following two subfamilies, the Terminal Uridyl Transferase (TUTase) and non-canonical Poly(A) Polymerase (ncPAP) ^13^. It has been reported that TUT4 (ZCCHC11) and TUT7 (ZCCHC6) are the major TUTases in mammals and are responsible for the oligo-uridylation at the 3’ ends of mRNAs, especially those with short A-tails (less than 25nt), which accelerates the degradation from both the 5’ and the 3’ ends of mRNAs ^7^. Some ncPAPs, such as TENT4A (PAPD7) and TENT4B (PAPD5), are able to incorporate guanyls amid A-tail extension in 3’ ends of mRNAs. This phenomenon, or mixed tailing, stabilizes poly(A) tails and increases the half-life of mRNAs ^9,14^.

The physiological significance of the TENT family has been revealed through generating mutant phenotypes. The function impairment of TUT4/7 and their orthologues lead to severe consequences including defects in early embryonic development, infertility and death in flies, zebrafish, *Xenopus*, and mice ^15-18^. TENT family enzymes also are active players in viral infection and host defense mechanisms. Cellular TENT4A/B can be hijacked by the hepatitis B virus (HBV), human cytomegalovirus (HCMV), and Hepatitis A virus (HAV) during infection in order to stabilize viral RNAs ^14,19,20^. Host cells also have defense mechanisms to combat viral infection, TUT4 and TUT7 are able to add oligo U tails to the virally derived RNAs to accelerate the degradation of viral RNAs during the infection of *C. elegans* by Orsay virus (OrV) and the infection of mammalian cells by influenza A virus (IAV) ^21,22^. Studies of TENTs hitherto have mainly uncovered their physiological functions in above processes.

Circadian clocks are timing devices that enable organisms such as animals, plants, fungus *Neurospora* and Cyanobacteria, to proactively adjust their physiology to daily oscillations according to the rotation of the earth. Endogenous, self-sustaining, and approximately 24 h length cycles generated by conserved cell autonomous transcriptional feedback loops have been a main focus of research for the past few decades ^23,24^. The main transcription/translation feedback loop (TTFL) in mammals includes two heterodimer complexes, one being a transcriptional activator complex CLOCK:BMAL1 and the other a transcriptional repressor complex PER:CRY ^25^. The CLOCK and BMAL1 heterodimer can transcriptionally activate the expression of clock-controlled genes including *period* (*per)* and *cryptochrome* (*cry)*. When the PER and CRY proteins are synthesized in the cytoplasm, they form heterodimers and are post-translationally phosphorylated by kinase CKIα, CKIδ and CKIε. Phosphorylation leads the translocation of PER:CRY heterodimers to the nucleus. There, they interact with CLOCK:BMAL1, which inhibits further transcription of clock-controlled genes ^26-28^. As PER and CRY proteins are degraded through the ubiquitin pathway, repression on CLOCK:BMAL1 is relieved and the cycle begins again with 24 h periodicity ^29,30^. Similarly, in flies, TTFL is also comprised of a transcriptional activator complex CLOCK:CYC, and a transcriptional inhibitor complex PER:TIM, regulation of the core circadian clock also shares resemblance ^31^. There is a secondary TTFL regulating the rhythmicity BMAL1 in mammals. In aammals, ROR and REV-ERB (NR1D) can be transcribed under CLOCK:BMAL1 activation, in turn, ROR activates the transcription of BMAL1, while REV-ERB blocks its transcription ^28^.

Although transcriptional regulation is the most important event in the rhythmicity, the post transcriptional steps including the processing, nuclear export, stability, and translation of mRNAs are also responsible for the maintenance of biological rhythms ^32-37^. Microarray analysis after fractionation of mRNAs according to their poly(A) tail lengths reveals that the rhythmic expression of cytoplasmic polyadenylation element-binding proteins (CPEBs) and the deadenylase Nocturnin (NOC) affects poly(A) tail lengths and translation of some mRNAs, suggesting that CPEBs and NOC are involved in the formation of Poly(A) Rhythmic (PAR) mRNAs ^35,37-39^.

The introduction and development of TAIL-seq technique has facilitated the probe into the 3’ ends of mRNAs, including 3’ end modifications and poly(A) tail lengths, globally and with increasing precision ^40^. TAIL-seq adopts a ligation-based method to prepare sequencing libraries and carries out paired-end sequencing to examine poly(A) tail regions and the corresponding gene regions. It processes the original sequencing signals with an improved base call algorithm so that poly(A) tail lengths can be correctly determined. Through the above procedures, TAIL-seq reveals poly(A) 3’ end modifications along with tail length data.

By sequencing the transcripts from mouse liver collected from 6 different time points in 12 h light:12 h dark (LD) daily cycles using TAIL-seq, we identified a subset of mRNAs with rhythmic poly(A) tail lengths in a daily dark-light circle. Furthermore, overall uridylation frequencies of 3’ ends of mRNA A-tails followed a circadian rhythm. However, guanylation or cytidylation did not exhibit these patterns. Consistent with the above results, major terminal uridylyl transferases, TUT4 and TUT7, showed cycling protein expression in mouse liver and the double knockdown of TUT4 and TUT7 in U2OS cells changed the rhythmic expression patterns of CLOCK genes. These results strongly suggest that rhythmic expression of TUTases contributes to the cycling of overall uridylation, and TUT4 and 7 play significant roles in mammalian circadian cycles.

## Results

### TAIL-seq identified the mRNAs with cycling poly(A) tail lengths and revealed the rhythmicity of overall 3’ terminal uridylation frequencies

To examine the diversity of mRNA poly(A) tail lengths and 3’ end modifications throughout the day, we performed TAIL-seq on LD-entrained mouse liver samples collected every 4 hours over 24 h period in 12 h light:12 h dark (LD) cycles. The representative clock genes Dbp and Rorc (NR1F3) showed rhythmic mRNA abundance as previously reported (Fig. S1A) ^41^. Rhythmicity of the poly(A) tail lengths and mRNA abundance of each gene were analyzed by Fourier Analysis (Fig. 1A) ^42,43^. This analysis identified 1032 genes with rhythmic poly(A) tail lengths, 1425 genes with rhythmic mRNA abundance, and 474 genes with both (Fig. 1A&1B &S1B, Suppl. table1&2). We observed that the poly(A) tail lengths of mRNAs negatively correlated with uridylation frequencies of mRNAs (Fig. 1B). The peaks of rhythmic A-tail lengths mostly appeared around ZT2 and ZT6 (Fig. 1C). The long poly(A) tail lengths were not in obvious accordance with high mRNA levels, probably because increased mRNA stability is not the only deciding factor in the steady state mRNA level (Fig. 1B) ^44^.

**Figure 1.**
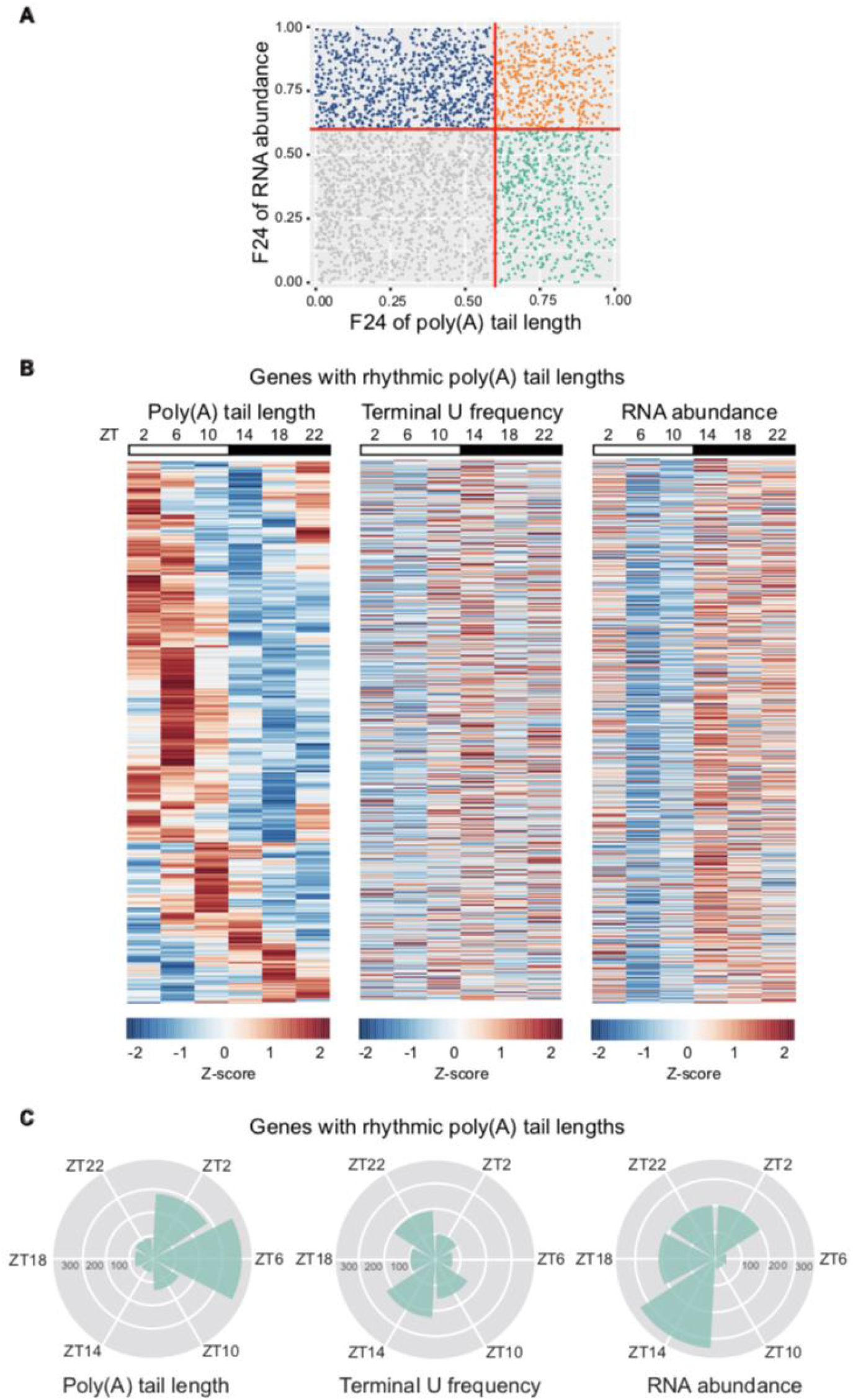
The identification of genes with rhythmic poly(A) tail lengths or rhythmic steady state mRNA expression by TAIL-seq. **(A)** The standardized RPMs and mean poly(A) tail lengths of each gene from 6 zeitgeber times (ZT) were analyzed by Fourier analysis. The 24 h spectral power F24 scores (range from 0 to 1) indicate the relative strength of the extracted circadian components. With a F24 score cutoff of 0.6 (red lines), 1032 genes display rhythmic poly(A) tail lengths (turquoise green and orange dots), 1425 genes are with rhythmic mRNA abundance (navy blue and orange dots), and 474 genes are with both rhythmic poly(A) tail lengths and mRNA abundance (orange dots). **(B)** The phase-sorted heatmap of identified 1032 genes with rhythmic poly(A) tail lengths from LD-entrained mouse liver. Each row represents one gene, separated by vertical panels in zeitgeber time. Z-scores from the standardized mean poly(A) tail lengths (left), 3’ terminal U frequencies (middle) and RPMs (right) of genes with rhythmic poly(A) tail lengths are shown. **(C)** Peak time distribution of mean poly(A) tail lengths (left), 3’ terminal U frequencies (middle) and RPMs (right) of genes with rhythmic poly(A) tail lengths. Each wedge represents a 4-h bin.

The top ten KEGG pathways enriched in the genes with rhythmic poly(A) tail lengths or rhythmic mRNA abundance overlapped in some fields, including thermogenesis, peroxisome, and valine, leucine and isoleucine degradation pathway. The pathways including cofactor biosynthesis, pyruvate metabolism, and beta-alanine metabolism were enriched only in the rhythmic poly(A) tail-length genes (Fig. 2A).

**Figure 2.**
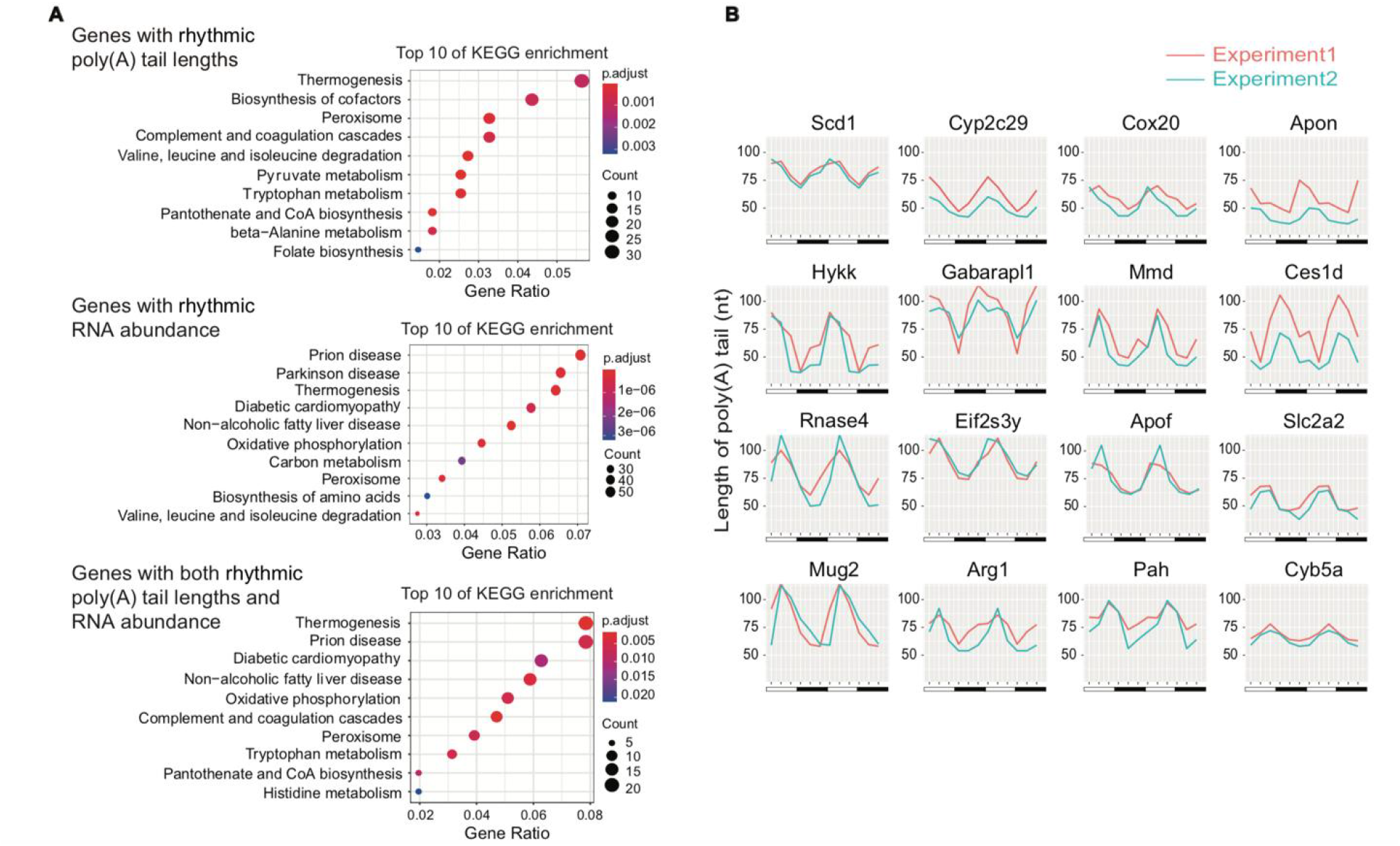
Functional enrichment analysis of rhythmically expressed genes. **(A)** KEGG enrichment analysis indicates the top ten enrolled pathways of the genes with rhythmic poly(A) tail lengths (the top panel), with rhythmic RNA expression (the middle panel), and with both (the bottom panel). **(B)** Representative oscillation of poly(A) tail lengths is shown. Red and blue lines represent poly(A) tail lengths from two independent experiments. The poly(A) tail lengths are double plotted for easier visualization of rhythmicity.

The liver is a metabolic center of carbohydrate, lipids, amino acids, and also the main organ controlling heat production. Peroxisome is the organelle where the reduction of reactive oxygen species (ROS), the catabolism of very long/branched chain fatty acids and bile acid intermediates occur. This suggests that rhythmic poly(A) tail lengths and mRNA levels contribute to animal metabolic regulation and thermogenesis adapting to food intake and environmental change. Some typical mRNAs with rhythmic poly(A) tail lengths include Eif2s3y, Eif4ebp3, Rnase4, Scd1, Cyp2c29, Cox20 et al. (Fig. 2B&S1C). The functions of Eif2s3y and Eif4ebp3 are related to translation initiation. Eif2s3y is a necessary for recruitment of initiator tRNAMet-tRNA^i^ to the 40S ribosomal subunit; while the Eif4ebp3 binds to Eif4e and prevents Eif4e from assembling into the translation initiation complex Eif4F. Pancreatic ribonuclease Rnase4 belongs to ribonuclease A superfamily and plays roles in mRNA cleavage, having specificity towards the 3’ side of uridine nucleotides (Fig. 2B).

We next analyzed the composition of 3’ ends of transcripts, that is, the frequencies of uridine, guanine and cytosine appearing at the 3’ ends of RNAs with different A-tail lengths. We found that the 3’ ends of transcripts with A-tails were often compromised with non-A nucleotides. Uridylation frequently appears in the transcripts with short A-tails (shorter than 25 nt), while guanylation often occurs in the transcripts with long A-tails (longer than 25 nt), consistent with previous observations (Fig. 3A and 3B) ^40^. Furthermore, we reproducibly observed that the frequencies of uridylation, especially oligo-uridylation (two or more uridines) fluctuated periodically throughout the day, having the lowest frequency around ZT2 and ZT6 (Fig. 3B and 3C).

**Figure 3.**
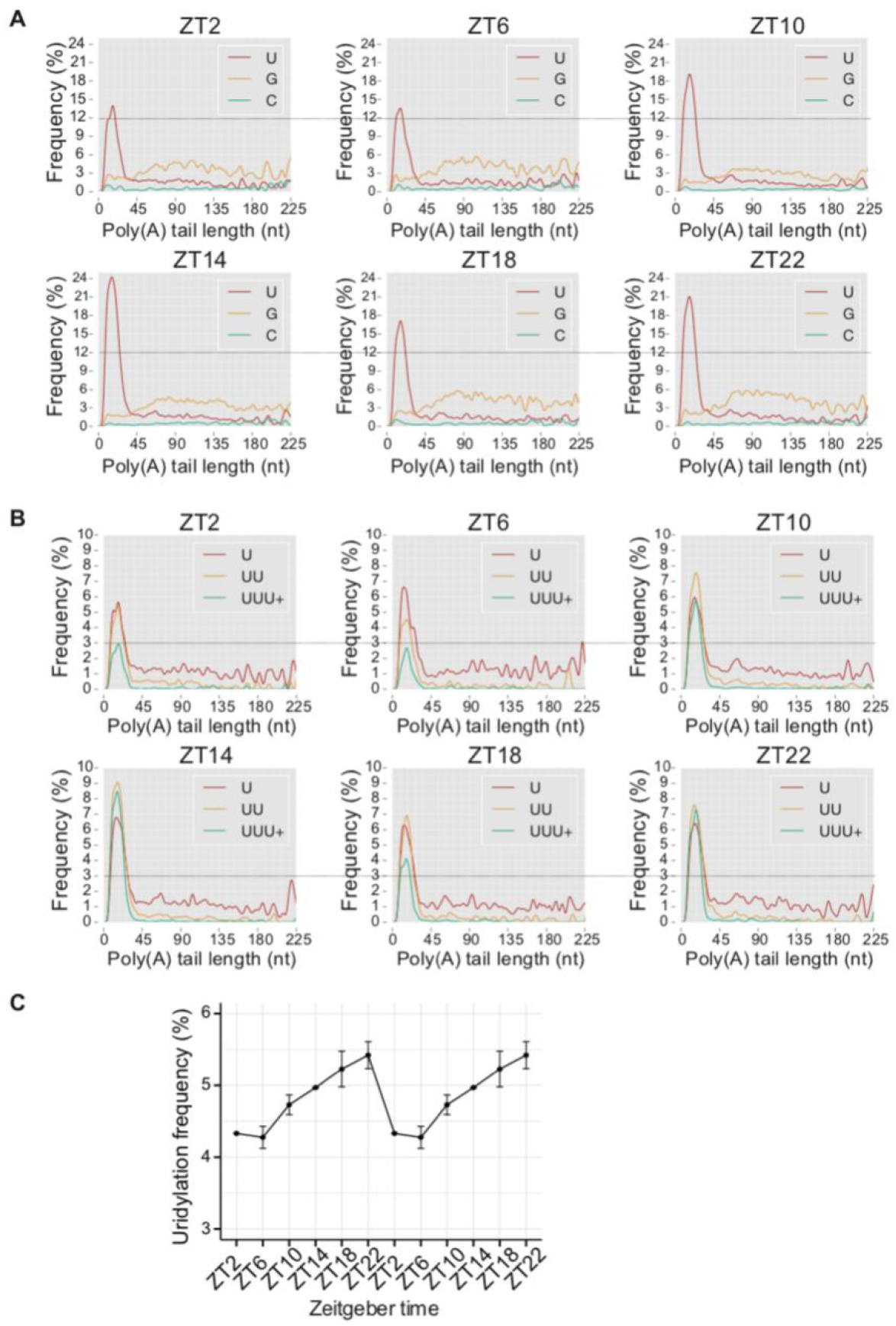
The uridylation frequencies at the 3’ ends of A-tails have a rhythmic change. **(A and B)** The distribution of modifications at the 3’ terminals of transcripts with different A-tail lengths shows that uridylation is enriched at the ends of transcripts with short A-tails (<25 nt), guanylation is enriched at the ends of transcripts with long A-tails (>25 nt), and cytidylation is only detected at low frequencies. The uridylation, guanylation and cytidylation frequencies (A), and mono-, di-, oligo-uridylation frequencies (B) from 6 zeitgeber times are shown. **(C)** The overall 3’ terminal uridylation frequencies of A-tailed transcripts are cycling in a daily dark-light circle (n=2, mean+/-SEM). The uridylation frequencies are double plotted for easier visualization of rhythmicity.

### Rhythmic expression of Terminal Uridyl Transferase TUT4 and TUT7

As described above, through performing TAIL-seq on LD-entrained mouse liver samples, we identified a subset of genes with cycling poly(A) tail lengths, where the wave pattern of dynamic A-tail lengths was in opposition to that of terminal uridylation frequencies. Deeper analysis pinpointed that overall uridylation frequencies oscillated and had the lowest value around ZT2 and ZT6. This hinted that some type of regulation exists in overall uridylation. So next we examined the expression of TUT4 and TUT7, which are mainly responsible for 3’ terminal uridylation in mammals. TUT7 showed rhythmic change in poly(A) tail lengths during 24 h period (Fig. 4A) and TUT4 mRNA had cycling steady state levels like reported (Fig. 4B&S1D) ^39,45^. Meanwhile, both TUT4 and TUT7 protein levels displayed a rhythmic pattern. We concurrently observed that total TUTase proteins maintained a relatively high expression level from ZT6 to ZT18 but made a sharp decrease at ZT22 (Fig. 4C&4D). Interestingly, consistent with the high protein expression of TUTase from ZT6 to ZT18 (Fig. 4D), cycling uridylation frequencies were high from ZT10 to ZT22 (Fig. 3C). The rhythmic oscillation on TUTase protein levels and uridylation frequencies displayed the phenomenon of time-delay, suggesting that rhythmic expression of TUT4 and TUT7 contributes to cycling uridylation frequencies in mouse liver.

**Figure 4.**
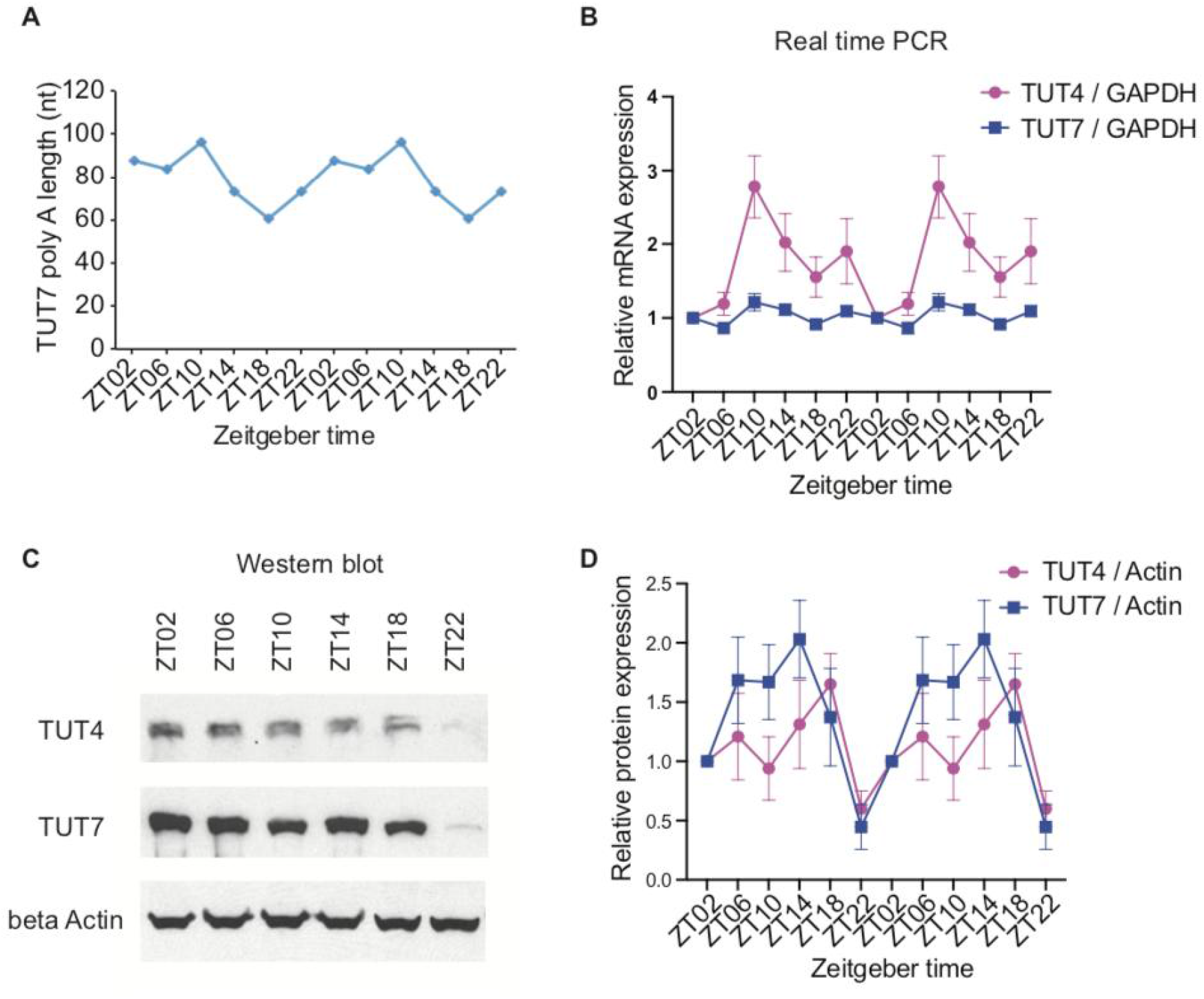
Rhythmic expression of TUT4 and TUT7. **(A)** poly(A) tail lengths of TUT7 from 6 zeitgeber times measured by Tail-seq in liver. **(B-D)** The mRNA levels (B) and protein levels (C-D) of TUT4 and TUT7 measured by real time PCR and western blotting respectively in liver (n=5, mean +/-SEM). The graphs are double plotted for easier visulization of rhythmicity.

### The contribution of TUTases to circadian expression of clock genes

It was well established in several organisms that oligo-uridylation on RNA triggers the downstream RNA decay ^46,47^. In the RNA decay pathway, TUT4/7-mediated mRNA 3’ terminal uridylation accelerates mRNA decay and shortens the half-life of mRNA in mammals ^7^. Therefore, we hypothesized that TUTases might contribute to the rhythmic expression of clock genes and influence molecular rhythm in a widespread and previously unrecognized manner. There have been no preceding reports that studied the functions of uridylation in biological rhythm to date. Remarkably, most tissues and some cells have autonomous circadian clock and the ability to sustain molecular oscillation in vitro ^48,49^.

To understand the function of TUTases in circadian oscillation, we subsequently simultaneously depleted TUT4 and TUT7 using RNAi in U2OS cells, one of the conventional models to investigate the cell autonomous oscillator ^50,51^. The depletion of TUT4 and TUT7 proteins were examined 48 h or 96 h following siRNA treatment, and were determined to be efficient through Western blotting (Fig. 5A). Following successful siRNA knockdown confirmation, TUT4 and TUT7 depleted U2OS cells were collected every 4 hours for 48 h, starting 25 h after cell synchronization by dexamethasone. The mRNAs of the core clock gene Per3 and NRID2 had rhythmic expression with a period length of 24 h in U2OS cells as detected by quantitative RT-PCR (Fig. 5B). Notably, the rhythmic amplitude of Per3 and NR1D2 mRNAs were dropped significantly in the groups of TUTase knockdown compared with control group (Fig. 5B and 5C). While the rhythmicity of Per3 and NR1D2 mRNAs had no significant differences between two groups as validated by Fourier analyses, in which the one that F24 score is closer to 1 has better rhythmicity (Fig. 5D).

**Figure 5.**
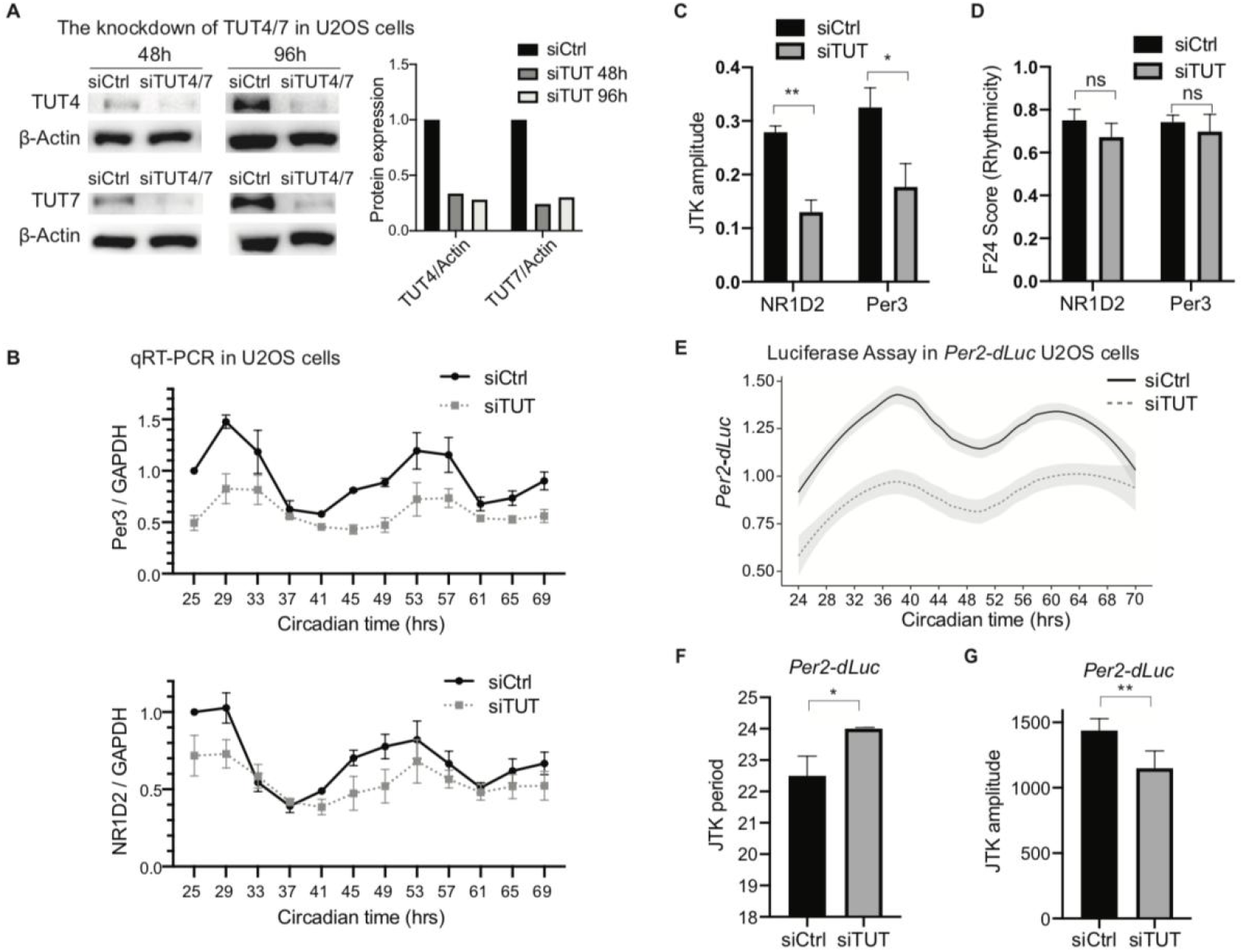
Knockdown of TUTases lengthens circadian period and reduces rhythmic amplitute of clock gene expression in U2OS cells. **(A)** U2OS cells were transfected with the mixture of siTUT4 and siTUT7 (siTUT4/7) or with siControl (siCtrl). The knockdown efficiencies of TUT4 and TUT7 were quantified by western blotting. The β-Actin was used as loading control. **(B-D)** After knockdown of TUT4/7 and cell synchronization, the Per3 and NR1D2 mRNAs in U2OS cells were measured by quantitative RT-PCR for 48 hrs in a 4-h bin and normalized to GAPDH (B). Their rhythmic amplitude from JTK cycle analysis (C) and F24 scores from Fourier Analysis (D) are shown (n=3, mean +/-SEM). P-value was calculated by a paired one-tailed *t* test. **(E-G)** Luciferase levels were measured in *Per2-dLuc* U2OS cells for 48 hrs in a 2-h bin after TUT4/7 depletion and cell synchronization (n=5, mean +/-SEM) (E). The rhythmic period (F) and amplitude (G) of luciferase expression analyzed by JTK cycle are shown. P-value was calculated by a paired one-tailed *t* test.

Next, the stable U2OS cell line of *Per2-dLuc* was used to monitor the rhythmic protein expression of Luciferase. In the *Per2-dLuc* cells, the gene transcription of rapid degradable luciferase protein was driven by the *per2* promoter. Interestingly, double knockdown of TUT4 and TUT7 extended the circadian period of luciferase in *Per2-dLuc* cells (Fig. 5E and 5F). Consistently, we noticed knockdown of TUTases reduced the cycling amplitude of luciferase (Fig. 5G). All these results indicate that mRNA terminal uridylation modulated by TUTases contributes to the rhythmic oscillation of clock genes.

In summary, we identified mRNAs with rhythmic steady state levels and/or poly(A) tail lengths and observed that overall frequencies of mRNA 3’terminal uridylation had a rhythmic change with the lowest value at ZT2 and ZT6. The protein levels of TUT4 and TUT7 were consistently low at ZT22 and the double knockdown of TUT4 and TUT7 reduced cycling amplitude and extended circadian period as explored by luciferase expression in *Per2-dLuc* cells.

## Discussion

The RNA decay pathway has been reported to participate in the circadian rhythm, such as the exosome in *Neurospora* and GW182 in *Drosophila* ^34,52^. Our study uncovered the key roles of uridylation, which triggers the RNA decay pathway, in circadian rhythm. We unveiled the delayed pattern between TUT4/TUT7 protein expression and overall mRNA 3’terminal uridylation frequencies in a 24 h dark-light cycle. As TUT4 and TUT7 are already established as the primary enzymes responsible for 3’terminal uridylation of mRNA, the controlled expression of TUTases will lead to overall change in mRNA uridylation. We also discovered that the TUTases are involved in the rhythmicity of clock molecules, revealed by the quantitative RT-PCR of Per3 and NR1D2 mRNAs and luciferase reporter assay of *Per2-dLuc*. It has yet been confirmed whether uridylation acts directly or indirectly on clock molecules, but it is clear that both uridylation and TUTases’ roles are critical for the molecular rhythmicity. Also, it is more likely that the uridylation affects overall mRNA metabolism but due to the importance of clock genes in circadian rhythmicity, the contribution of uridylation on some clock genes may be more significant for cellular autonomous clock formation.

The dynamic change in overall uridylation frequencies has been detected in early embryonic developmental stages of vertebrate and in different phages of the mammalian cell cycle ^16,53^. Interestingly, in zebrafish and *Xenopus*, the differences in TUTase expression also cause the differences in uridylation frequencies in early developmental stages, not unlike our observation in the mouse liver. In early embryonic development, zygotic genome activation results in the expression of TUT4/7 and further leads to increased uridylation of maternal mRNAs, triggering the swift removal of maternal transcripts ^16^.

Several poly(A) tail detection methods have been developed so far, such as TAIL-seq, mTAIL-seq, PAL-seq, PAL-seq V2, PAT-seq, TED-seq, FLAM-seq, PAIso-seq and other emerging methods ^3,5,11,12,40,54-57^. Along with advances in A-tail assessment strategies, the functions of 3’ terminal modifications in key physiological processes have been elucidated, including fertility, early embryonic development, viral infection, and host defense in *C. elegans*, flies, zebrafish, *Xenopus*, and mammals ^47,58^. Here, we have manifested for better understanding, the impact of dynamic change and the function of uridylation regarding the circadian pathway. It will be fascinating to further examine the dynamic changes of mRNA 3’terminal modifications and their functions in varying species and biological processes and illustrate the underlying mechanisms.

### EXPERIMENTAL PROCEDURES

#### Library preparation and analysis of TAIL-seq

Library preparation and analysis of TAIL-seq were conducted according to protocol described by Chang with minor adjustments ^40^. Male C57BL/6 mice were housed under 12 h light:12 h dark (LD12:12) with food and water available ad libitum, and tissues were collected in 3-5 month-old mice. All animals were used in accordance with the guidelines set forth by the Institutional Animal Care and Use Committee of Texas A&M University. Total RNA was extracted from LD-entrained mouse liver samples collected every 4 hours over 24 h in a 12 h light:12 h dark (LD) cycle. TAIL-seq libraries were sequenced by Illumina Hiseq 2500 (51 × 251 bp paired end run) mixed with 20% PhiX control library to balance base distribution, and 1% spike-ins with standard lengths of poly(A) tails as internal reference. Then, read 1 (51 bp in length) of the tags were aligned against UCSC mm10 and original fluorescence signals of read 2 (251 bp in length) were used for poly(A) length calculation with the base-call strategy. More details are included in the supplemental experimental procedures.

#### Functional enrichment analysis

Kyoto Encyclopedia of Genes and Genomes (KEGG) pathway enrichment analysis was carried out using the clusterProfiler package (V3.118.1) ^59^. Before the analysis, we used g:Profiler website ^60^ to convert gene symbols to Entrez ID. KEGG pathways with p-value < 0.05 were considered as significantly enriched pathways. The top 10 of KEGG pathways were shown.

#### *Per2-dLuc* U2OS stable cell culture

U2OS cells stably expressing *Per2-dLuc* reporter (a generous gift from John Hogenesch) were maintained in DMEM medium containing 10% fetal bovine serum (FBS), 2mM L-Glutamine (Invitrogen), 0.1 mM nonessential amino acids (Invitrogen) and 1×P/S (Invitrogen) at 37 degrees in a humidified incubator supplied with 5% CO2 ^50^. The *dLuc* gene was driven by the *per2* promoter including 526 bp upstream of the *per* transcription start site. The *dLuc* construct contains a *luciferase* gene and a PEST sequence for rapid protein degradation as previously described ^61^.

#### siRNA transfection and rhythmicity assay in U2OS cells

The rhythmicity assay conducted in *Per2-dLuc* U2OS cells followed the protocols described for quantitative RT-PCR ^51^ and for luciferase assay ^50^ with minor adjustments. On the first day (Day 1), *Per2-dLuc* U2OS stable cells were seeded in 12-well plate (Luciferase assay) or 6-well plate (quantitative RT-PCR). On the second day (Day 2) and the fourth day (Day 4), cells were transfected with siRNA mixture against TUTases (Sigma, 50 nM of final concentration) using Lipofectamine 2000 transfection reagent (Invitrogen) following the manufacturer’s instructions. A negative control siRNA (Sigma) was used to ensure that equal molar amounts of siRNA was added in each condition. Eight hours after the second transfection of siRNA (Day 4), the cells were synchronized by 0.1 μM dexamethasone (Solarbio), and collected every 2 hours (Luciferase assay) or 4 hours (quantitative RT-PCR) for 48 h, starting 1 day after cell synchronization (Day 5). For Luciferase assay, luminescence levels were measured using Dual-Luciferase® Reporter Assay System (Promega) and Cytation™ 3 Cell Imaging Multi-Mode Reader (BioTek). Real time PCR of NR1D2 and Per3 were conducted using TB Green Premix Ex Taq kit (Takara) in Q3 real-time PCR system (Applied Biosystems).

The following siRNAs were used: siCtrl-S, 5’-CCU ACG CCA CCA AUU UCG UUU-3’; siCtrl-AS, 5’-ACG AAA UUG GUG GCG UAG GUU-3’; siTUT4-S, 5’-GGA AUG AAG AAG AGA AAG AUU-3’; siTUT4-AS, 5’-UCU UUC UCU UCU UCA UUC CUU-3’; siTUT7-S, 5’-GAA AAG AGG CAC AAG AAA AUU-3’; siTUT7-AS, 5’-UUU UCU UGU GCC UCU UUU CUU-3’. The following primers were used: Per3-F, 5’-CCA GAC TGT CAC TCA AGA AAT-3’, Per3-R, 5’-CAG CTG TCT TCT ACC AGA AC-3’; NR1D2-F, 5’-GGA CAT GAA ATC TGG GAA GAA-3’, NR1D2-R, 5’-AGA TCT CTG AAC CCA GGA ATA-3’.

#### Cycling analysis using JTK cycle and Fourier transformation

The mRNAs with cycling steady state mRNA expression or with rhythmic poly(A) tail length were identified using Fourier transformation ^43^. To be considered as cycling, the following cutoff thresholds were used: a F24 score of greater than 0.6, and p-value of less than 0.05, mean poly(A) tag counts of greater than 30, a mRNA cycling amplitude (maximum expression divided by minimum expression) of at least 1.5-fold, a cycling poly(A) tail length amplitude (maximum A-tail length minus minimum A-tail length) of at least 10 nt.

Both Fourier transformation ^43^ and the JTK cycle component of MetaCycle ^62^ were used for the cycling analysis of Per3 and NR1D2 mRNAs in U2OS cells. Each replicate of quantitative RT-PCR data was separately analyzed by Fourier transformation and JTK cycle. Then, the average value of F24 scores and JTK amplitudes were calculated from replicates. P-value was calculated by a paired one-tailed *t* test.

The JTK cycle was used for the cycling analysis of *Per2-Luc* luminescence signals in U2OS cells. The average of luciferase assay from each two replicates was treated as a set and each set of data was used for JTK cycle analysis. Then the mean and p-value (a paired one-tailed *t* test) of JTK period and JTK amplitude from replicate sets were examined. To plot the graph of *Per2-Luc* luminescence signals against circadian times, the mean and SEM from 5 replicates of experiments were used. The graph was fitted using geom smooth function of ggplot2 package in RStudio.

## Supporting information

Supplemental Experimental Procedures

## Supplementary materials

Supplemental table 1: Genes with rhythmic poly(A) tail lengths

Supplemental table 2: Genes with rhythmic mRNA expression

Supplemental Experimental Procedures

## Acknowledgements

We gratefully acknowledge Michael Rosbash and his lab for having hosted Hua Jin during the beginning of the work. We thank John Hogenesch for U2OS luciferase reporter (*Per2-dLuc*) stable cell lines.

## Author contributions

H.J. is responsible for conceptualization. H.J., J.W., W.P., Y.Y., X.W., H.Z., C.L. and D.N. performed the experiments. J.M. provided mouse liver and RNA samples, W.S., H.J., A.S., and H.C. analyzed and visualized the data. H.J., W.S., Y.Y., J.W. and W.P. wrote the original draft. Y.L. and V.K. reviewed and edited the manuscript. H.J. is contributed to funding acquisition.

## Data and materials availability

All data needed to evaluate the conclusions in the paper are present in the paper and/or the Supplementary Materials. Raw next-generation sequencing data from this study were deposited at National Center for Biotechnology Information Sequence Read Archive database with accession number PRJNA898115. Additional data related to this paper may be requested from the authors.

---

**Supplementary Fig. 1.**
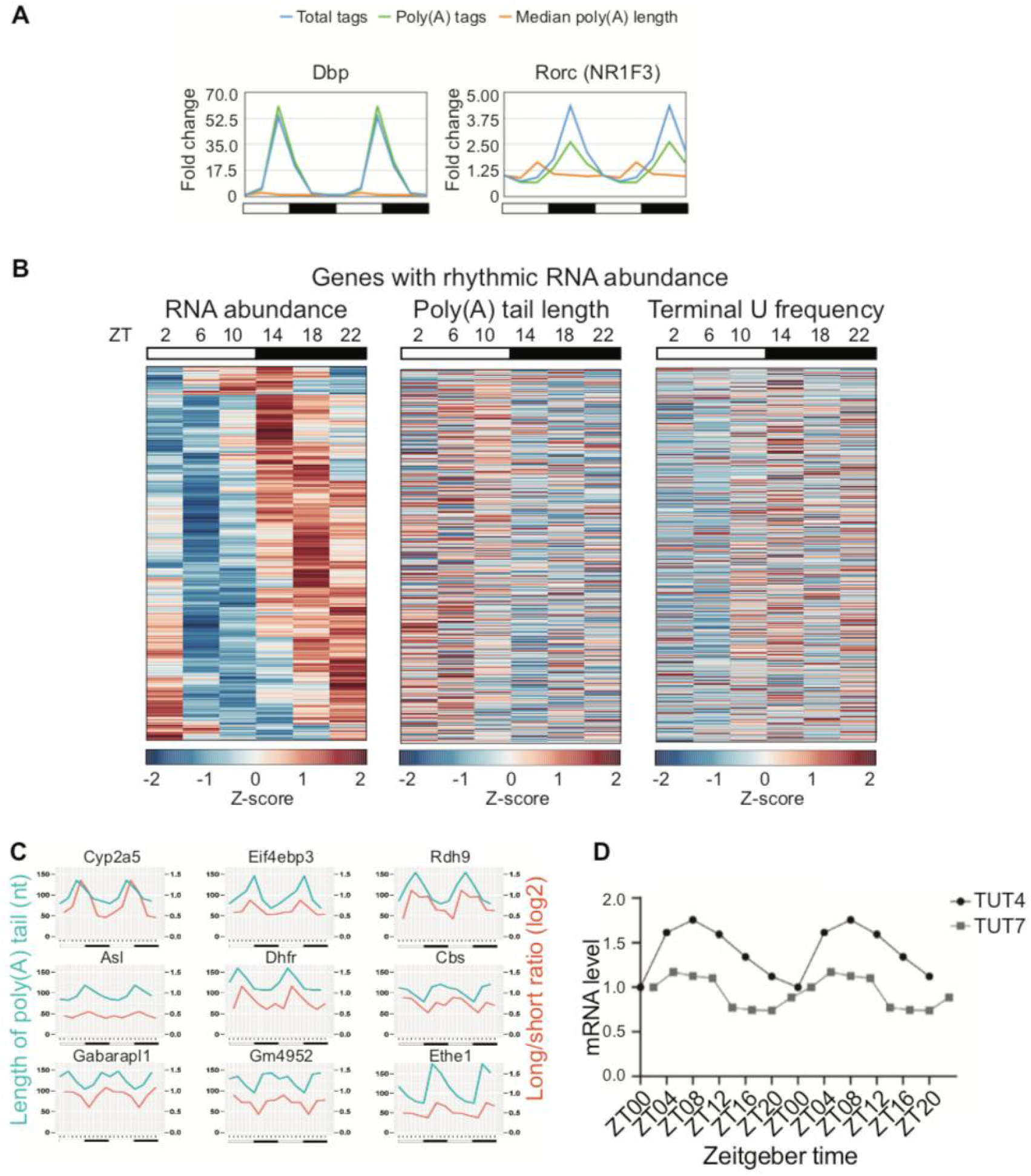
The genes with rhythmic mRNA abundance. **(A)** The mRNA oscillation of representative clock gene Dbp and Rorc (NR1F3) in mouse liver TAIL-seq are shown. **(B)** The phase-sorted heatmap of 1425 genes with rhythmic mRNA abundance identified from LD-entrained mouse liver. Each row represents one gene, separated by vertical panels in zeitgeber time. Z-score indicates the standardized values of RPMs (left), mean A-tail lengths (middle), and 3’ terminal uridylation frequencies (right) respectively. **(C)** Representative genes have similar oscillation of A-tail lengths as that in Kojima’s paper. The blue lines represent results from our Tail-seq data, and the red lines represent data from Kojima’s paper. All graphs are double plotted for easier visualization of rhythmicity. **(D)** The change in mRNA levels of TUT4 (microarray data from Kojima’s paper) and TUT7 (RNA-seq data from Yoshitane’s paper) are shown.

