## Supplemental Experimental Procedures for "The Role of Terminal Uridyl Transferases in the Circadian Rhythm"

### TAIL-seq library preparation and analysis

TAIL-seq library preparation and analysis were followed the guideline described in the paper (Chang et al., 2014) and the experiment was carried out in Brandeis University, USA. Total RNAs were extracted using TRIZOL reagent (Invitrogen) from LD-entrained mouse liver samples collected every 4 hours over 24-h period in 12-h light:12-h dark (LD) cycles. The total RNAs were treated with DNase I (Takara) and then size selected (>200nt) using RNeasy MinElute Clean up Kit (Qiagen). The rRNAs were removed using Ribo-zero Kit (Epicentre). The rRNA-depleted RNAs were ligated to 3' biotin-labeled DNA adapters (IDT synthesized), then partially digested with RNase T1 (Ambion). The digested RNAs were pulled down by Streptavidin beads (Invitrogen, 11206D), phosphorylated the 5' end with PNK (Takara, 2021B), and size selected again (500-1000 nt) by gel purification. The purified RNAs were ligated to 5' adapters (IDT synthesized), reverse transcribed by Superscript III reverse-transcriptase (Invitrogen, 18080085), and PCR amplified by illumina True-seq PCR primers and Phusion polymerase (Thermo Fisher). The PCR products larger than 450 bp were purified using AMPure XP beads.

To prepare spike-in references, synthesized DNA fragments (IDT) with polyA tail lengths of 8 nt, 16 nt, 32 nt, 64 nt, 118 nt, and 128 nt were amplified respectively by 15 cycles of PCR using illumine True-seq PCR primers. PCR products were purified using PAGE gel, and quantified by Agilent Bioanalyzer. Each spike-in was mixed in the same mole ratio. Then, TAIL-seq libraries were combined with 20% PhiX control library (GenBank accession J02482.1) as well as 1% spike-in internal references, and sequenced by Illumina Hiseq 2500 (51 bp X 251 bp paired end run). The following oligos were used: RNA 5' Adapter (RA5), 5'-GUU CAG AGU UCU ACA GUC CGA CGA UC-3'; RNA 3' Adapter (RA3 biotin 4), /5App/CTG CAN NNN NNN NNN NNN NNT GGA ATT CTC GGG TGC

CAA GGC/iBiodT//iBiodT//3ddC/; RNA RT Primer (RTP), 5'-GCC TTG GCA CCC GAG AAT TCC A-3'; RNA PCR Primer (RP1), 5'-AAT GAT ACG GCG ACC ACC GAG ATC TAC ACG TTC AGA GTT CTA CAG TCC GA-3'; RNA PCR Primer, Index 1 (RPI1), 5'-CAA GCA GAA GAC GGC ATA CGA GAT CGT GAT GTG A CTGGAG TTC CTT GGC ACC CGA GAA TTC CA-3'.

For TAIL-seq data analysis, it was filtered out the read pairs that can be aligned to the common contaminant set including rDNA, PhiX genome, PCR primers, 5S and 5.8S rRNA sequences. Read 1 (51 bp) was aligned against UCSC mm10 genome. Original fluorescence signals of read 2 (251 bp) were used for polyA length calculation with the base-call strategy. A weight is provided to each base between "start position" to the "end position", under the condition that the sum of these base weights was above zero, the max length between the "start position" and the "end position" was identified as the primary length of the poly(A) tail. If the measured tail lengths were under 8 bp, the primary lengths were their real lengths. For the others, the real tail lengths were determined by secondary calibration by Gaussian-mixture-hidden-Markov-Model whose parameters were calculated from 500 random clusters of the spike-ins.

**Examination of nontemplated terminal nucleotidyl addition**

The analysis of nontemplated terminal nucleotidyl addition was followed the guideline described in the paper (Lim et al., 2014). To calculate uridylation frequency, very short poly(A) tails between 1 and 4 nucleotides were excluded. In addition, noncoding RNAs (mainly snoRNA precursors) and histone mRNAs were excluded as well since their metabolism were different from the major mRNAs. For shorter poly(A) tails (shorter than 10 nt), we filtered them again to eliminate artifacts from unwanted RNA ligation between the 3' end of an mRNA and the 5' end of abundant small RNAs (such as 5S rRNA, 5.8S rRNA, let-7, and miR-21). They were considered as poly(A) tails only if 80% or more in the nontemplated addition is composed of adenosines after trimming all consecutive nonadenosine nucleotides from the 3' ends. The lengths of U tails were measured by counting perfectly consecutive uridines from the 3' ends within the nontemplated addition. The frequency of terminal guanylation, cytidylation, or uridylation (mono-, di-, oligo-) was profiled in mRNAs with different A-tail lengths.

**Quantitative real-time PCR and western blot analysis in mouse liver**

For real-time PCR, total RNAs were purified as in the TAIL-seq. Reverse transcription was performed using 1.5 μg of RNA, M-MLV Reverse transcriptase (Promega) and oligo dT mixture (IDT synthesized). Real-time PCR was conducted using TB Green Premix Ex Taq kit (Takara) in Q3 real-time PCR system (Applied Biosystems). The mRNA abundance of each transcript was normalized to that of GAPDH obtained in the same cDNAs. Real-time PCR primers were designed on IDT website and used as follows: TUT4-F, 5'-CAG AGA ATG GTT GAT GGA TGG-3', TUT4-R 5'-CCA GTG ATT CCG TGT TCT TC-3'; TUT7-F, 5'-CGT GAT CAG CAT CAG AAG AA-3', TUT7-R, 5'-CTC GAT AAT CCA GCA CCA AG-3'; GAPDH-F, 5'-GGA GAA ACC TGC CAA GTA TG-3', GAPDH-R, 5'-AAC CTG GTC CTC AGT GTA G-3'.

For western blot analysis, mouse liver samples were collected as in the TAIL-seq and grinded respectively in liquid nitrogen. The grinded samples were lysed in RIPA buffer supplemented with 1mM phenylmethylsulfonyl fluoride (PMSF) for 30 minutes at 4 degrees. The supernatant was separated using 8% polyacrylamide SDS-PAGE gels, and wet-transferred to nitrocellulose membranes (GE Amersham). Then, the membranes were blocked with 4% milk (Fisher Sci Co.) in TBST for 30 min, then immunoblotted overnight at 4 degrees with primary antibody against TUT4 (1:500 dilution in 5% milk, from V. Narry Kim's lab), TUT7 (1:500 dilution in 1% milk, from V. Narry Kim's lab), or beta Actin (1:300 dilution in 4% milk, Santa Cruz Biotechnology). The membranes were washed three times with TBST buffer and incubated with horseradish peroxidase-conjugated goat anti-mouse/rabbit IgG (1:20000 dilution, Quality Yard) for 1 hour. After TBST washing, the membranes were incubated with enhanced chemiluminescence kit (Tanon ECL kit) right before the exposure on film or the measurement in iBright photographer (ThermoFisher). The results were qualified by ImageJ using beta Actin as the internal reference.
